# Incoherent modulation of bi-stable dynamics orchestrates the Mushroom and Isola bifurcations

**DOI:** 10.1101/2021.04.22.440901

**Authors:** Amitava Giri, Sandip Kar

## Abstract

In biological networks, steady state dynamics of cell-fate regulatory genes often exhibit Mushroom and Isola kind of bifurcations. How these complex bifurcations emerge for these complex networks, and what are the minimal network structures that can generate these bifurcations, remain elusive. Herein, by employing Waddington’s landscape theory and bifurcation analysis, we have shown that both Mushroom and Isola bifurcations can be realized with four minimal network motifs that are constituted by combining positive feedback motifs with different types of incoherent feedback motifs. Our study demonstrates that the intrinsic bi-stable dynamics due to the presence of the positive feedback motif can be fine-tuned by altering the extent of the incoherence of these proposed minimal networks to orchestrate these complex bifurcations. These modeling insights will be useful in identifying and analyzing possible network motifs that may give rise to either Mushroom or Isola bifurcation in other biological systems.

## Introduction

Biochemical networks are inherently complex and consist of several nonlinear molecular interactions(Kitano, 2002). However, in many biological processes, the cell fate decision-making events are organized by maintaining the steady-state level of an important regulatory gene in a bi-stable manner(Yao *et al.*, 2008; Sokolik *et al.*, 2015; Sengupta, Kompella, *et al.*, 2018; Sengupta, Govindaraj, *et al.*, 2018; TYSON and NOVAK, 2001) as a function of a specific external or internal signal. Recent studies further revealed that in the context of stem cell differentiation regulation, the related biological decision-making happens via more complicated dynamical processes such as Mushroom and Isola kind of bifurcations(Sengupta and Kar, 2018; Chickarmane and Peterson, 2008). Mushroom bifurcation even explains the biochemical mechanisms underlying synaptic strengthening and memory formation between organisms(Song *et al.*, 2006). Importantly, for both Mushroom and Isola bifurcations, two bi-stable switches get coupled in a particular way to create a definitive dynamical response for the corresponding biological network. However, little is known about the minimal network structure to dynamically observe such kind of bifurcation features, which seem to influence important regulatory events in biology.

Biological networks normally contain certain repetitive sub-networks, which are known as network motif(Alon, 2007). These network motifs are capable of producing specific dynamical responses. For example; a biochemical network can display bi-stable dynamics due to either auto positive regulation, double-positive feedback, or double-negative feedback motif(Kashtan *et al.*, 2004; Tyson *et al.*, 2003; Milo *et al.*, 2011). Likewise, feedforward network motifs in the form of incoherent feedforward loops are also well-known for showing adaptation behavior in many biological systems(Kaplan *et al.*, 2008; Wang *et al.*, 2020; Qiao *et al.*, 2019; Ma *et al.*, 2009). It appears that biological networks are pre-designed by combining these network motifs to give rise to complex dynamical responses such as tri-stability, Mushroom, and Isola bifurcation. Herein, we wanted to investigate the minimal network architectures that can give rise to either Mushroom or Isola bifurcation features as a function of any external signal. Knowing the basic design-principle of these biological networks will be crucial in understanding and eventually altering the dynamics of the various cell-fate regulatory developmental events that are governed by these novel bifurcations.

In this regard, existing mathematical models suggest that a positive feedback network motif is essential to produce a bi-stable system(Tyson *et al.*, 2003), which can be further modulated to generate a tri-stable, Mushroom, or Isola bifurcations by introducing other network motifs. It has been demonstrated that adding further positive feedback motifs to an existing bi-stable system can lead to tri-stable bifurcation scenarios in a biological system(Dey and Barik, 2021; Rata *et al.*, 2018; Tian *et al.*, 2013). On the contrary, models related to stem cell developmental dynamics indicate that an incoherent feedforward loop (IFFL) might be essential in addition to a positive feedback motif to create either Mushroom or Isola type bifurcations(Sengupta and Kar, 2018; Chickarmane and Peterson, 2008). However, it was not conclusive that these two motifs are enough to create these bifurcations or not, as other network motifs are also present in those complicated networks. Can it be generalized that incoherent feedback motifs in combination with a positive feedback regulation are sufficient to produce Mushroom or Isola bifurcations?

To explore this important question, we proposed a set of four networks (**Fig.1**) consisting of three components (*X, Y*, and *Z*), where we assumed that *Z* gene can activate its transcription via auto-positive feedback regulation. Additionally, we consider that the external signal (*S*) activates the transcription of the gene *X*, and *X* either activates or inhibits the formation of *Y* and *Z* in all possible combinations to give rise to four network configurations (**Fig.1**). The dynamical equations for the network components (**Table-S1**) are constructed by using mass-action kinetics as well as phenomenological terms to represent the positive feedback regulation and incoherence of the networks adequately (see **Supplementary text** for details). **Table-S2** contains all the values of kinetic parameters. Herein, we investigated two important questions. First, among these four networks, which are capable of producing Mushroom or Isola bifurcations? Second, if these combined network motifs do produce these unique bifurcations, then what factors determine the nature of the bifurcations and how to alter them to affect the response output in *Z*?

**Fig.1:**
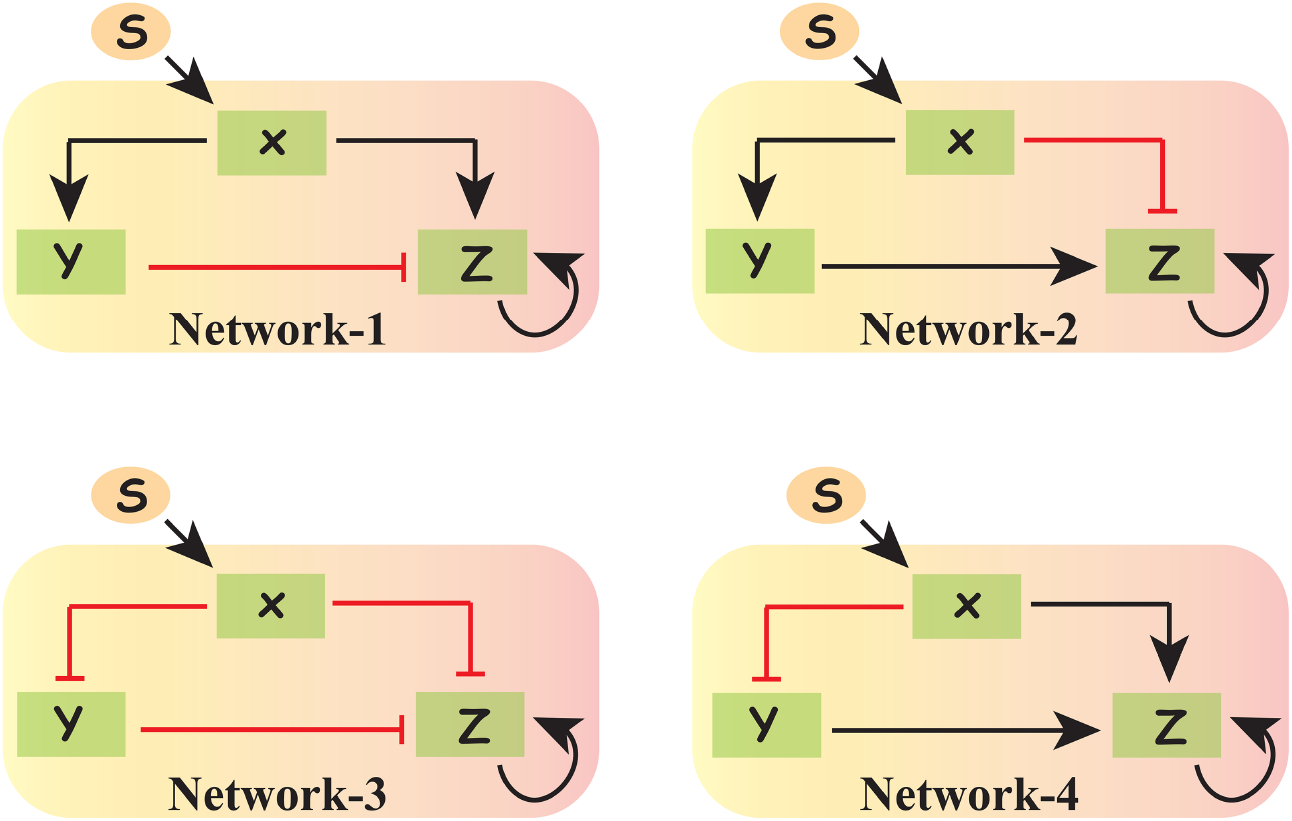
Four proposed network motifs that have the potential to produce Mushroom and Isola bifurcations. Red hammer headed arrows and black arrows represent inhibition and activation respectively.

## Results

### Potential landscape approach indicates the minimal network motifs for mushroom bifurcation

Recent studies have demonstrated that Waddington’s potential landscape theory can be considered as an alternative and simpler way to identify network motifs behind different cellular fate-determining events(Wang *et al.*, 2010; Ferrell, 2012). In a potential energy diagram, the local minima characterize the stable steady-state and the local maxima represent the unstable steady state(Ferrell, 2012). The multidimensional nature of the complex biological networks often leads to non-Newtonian dynamics, and calculating the potential energy for the respective biochemical network becomes extremely challenging. However, the simplicity of our proposed mathematical models allowed us to obtain the effective potential even though the systems are multidimensional. We calculated the effective force of the system by using the composite function method, where a function of several variables can be expressed into a single variable function(Dey and Barik, 2021). The expression for the effective force comes out as,

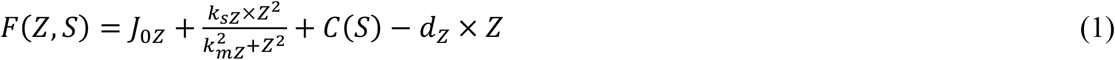
where 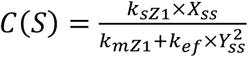 and *X_SS_, Y_SS_* are the steady-state expressions for the *X* and *Y*, respectively.

The potential *V*(*Z,S*) is defined as 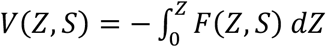, where *V*(*Z,S*) is a function of a single variable *Z.* The effective potential is a parametric function of the external signal (*S*). We further obtained the analytical expression of the effective potential as,

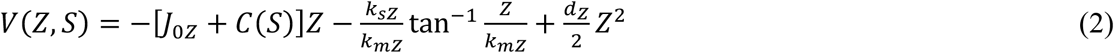

In **Fig.2a**, the potential is plotted as a function of *Z* for the Network-1 (**Fig.1**), which shows that the potential landscape changes its nature as the signal strength (*S*) is increased systematically. For *S* =5, there is only one local minimum for the lower value of *Z*, suggesting the existence of a single stable steady state. However, at *S* =10, the potential energy landscape has two local minima separated by a local maximum, which indicates that there are two stable and one unstable steady state. For *S* =17, the local minimum at higher values of *Z* becomes more stable, while the other local minima at lower values of *Z* disappeared. Intriguingly, at *S* = 30, the potential landscape regains two local minima separated by a local maximum indicating that the bi-stability in the network dynamics is restored (**Fig.2a**). For, higher signal strength (*S* = 45), the potential energy has only one local minimum reflecting a single stable steady state. We have monitored the changes in the values of the extrema according to their types for all values of *S*, and found that it indicates a Mushroom bifurcation (**Fig.2b**). Interestingly, the bifurcation diagram (**Fig.2c**) of *Z* as a function of *S* turns out to be identical to that obtained from the potential energy-based analysis (**Fig.2b**). In **Fig.2c**, the bifurcation diagram has been overlaid on top of the 2D-contour of the potential energy landscape for network-1 to demonstrate the correlation between the two different methods to decipher the dynamical presence of the mushroom bifurcation, where the points designated by 1-4 (**Fig.2c**) are the respective saddle nodes. Importantly, similar potential energy landscape analysis with network-2 (**Fig.S1a**) and network-3 (**Fig.S1b**) also produces the mushroom bifurcations. However, the relative stabilities of the individual minima varied for different networks (**Fig.S1**), which eventually altered the extent of the bi-stable regions and the shapes of the mushroom bifurcations (**Fig.2d** and **Fig.2e**).

**Fig.2:**
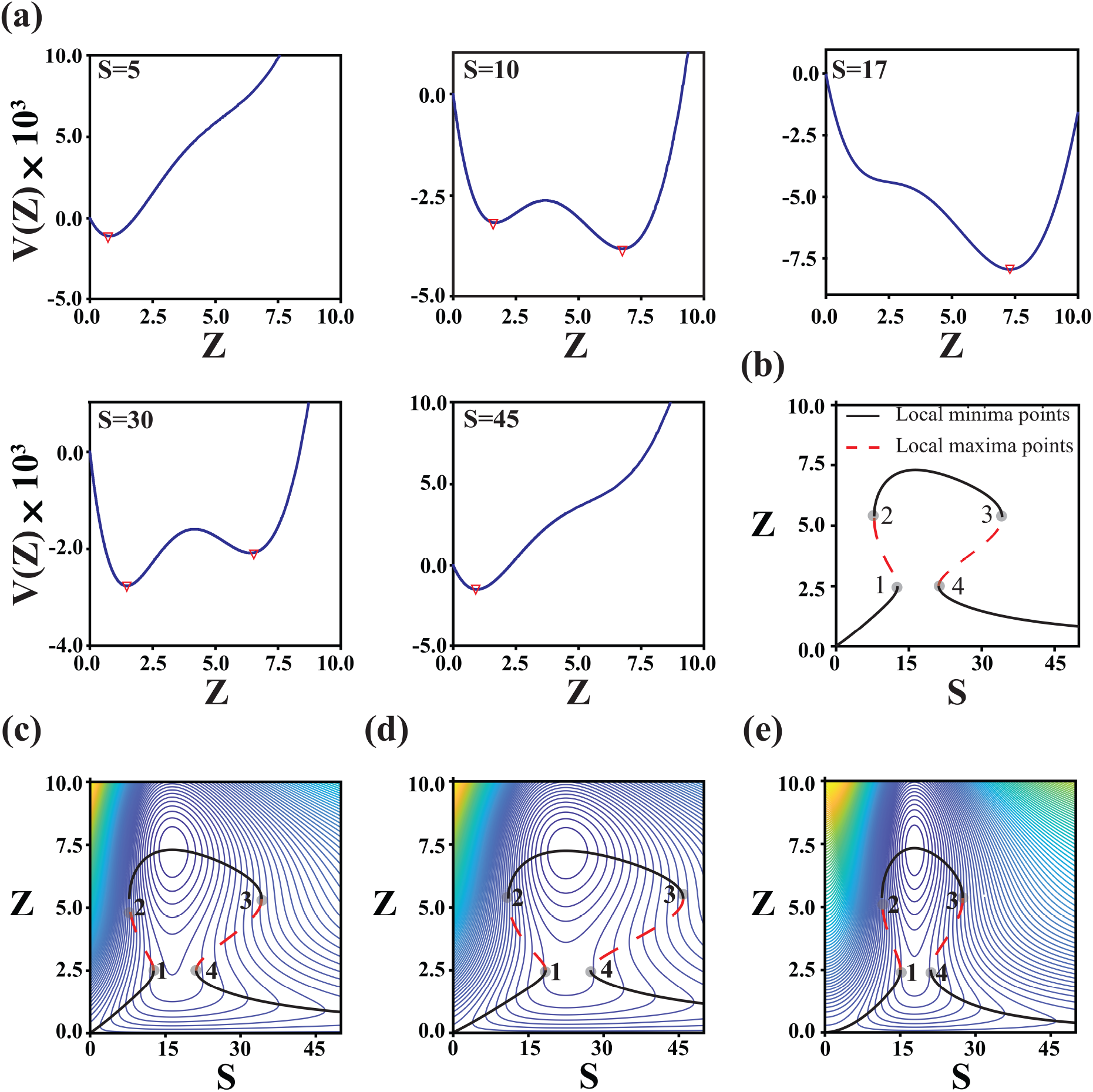
Potential landscape analysis reveals the existence of mushroom bifurcation in the proposed networks. **(a)** The plot of effective potential as a function of *Z* for different values of *S*. **(b)** Energy-based bifurcation diagram of the steady state of *Z* as a function of *S* for network-1. Eigen value-based bifurcation diagrams of *Z* Vs. *S* is overlayed on top of the contours of the potential energy landscapes for **(c)** Network-1, **(d)** Network-2, and **(e)** Network-3.

### Isola bifurcations dynamically emerge from the same combined network motifs

From the perspective of bifurcation theory, it is known that a mushroom bifurcation can lead to Isola bifurcation if the saddle nodes SN_1_ and SN_4_ (**Fig.2c-e**) dynamically converge and annihilate each other. In a multi-parametric system, this can be achieved in several ways. However, in our simple models (**Table-S1**), the extent of positive feedback and the degree of incoherence is mainly responsible for the dynamical diversity of these networks. On reducing the positive feedback regulation (for Network-1), the potential energy landscapes for *S* = 5, 10, 30, and 45 manifest only a single local minimum (**Fig.3a**) corresponding to a low level of *Z* However, for *S* = 17, the potential energy landscape contains two local minima and one local maximum, which represents a bi-stable scenario and dynamically is in stark contrast to the observations made previously (**Fig.2c**). The energy-based 1-p bifurcation diagram as a function of *S* demonstrates the occurrence of an Isola bifurcation (**Fig.S4a**). The eigenvalue-based 1-p bifurcation plotted on top of the 2-D contour (**Fig.3b**) of the corresponding potential energy landscape further reconciles the energy-based bifurcation and illustrates the relative positions of the saddle nodes (2 and 3) indicating the relative stabilities of the steady states.

**Fig.3:**
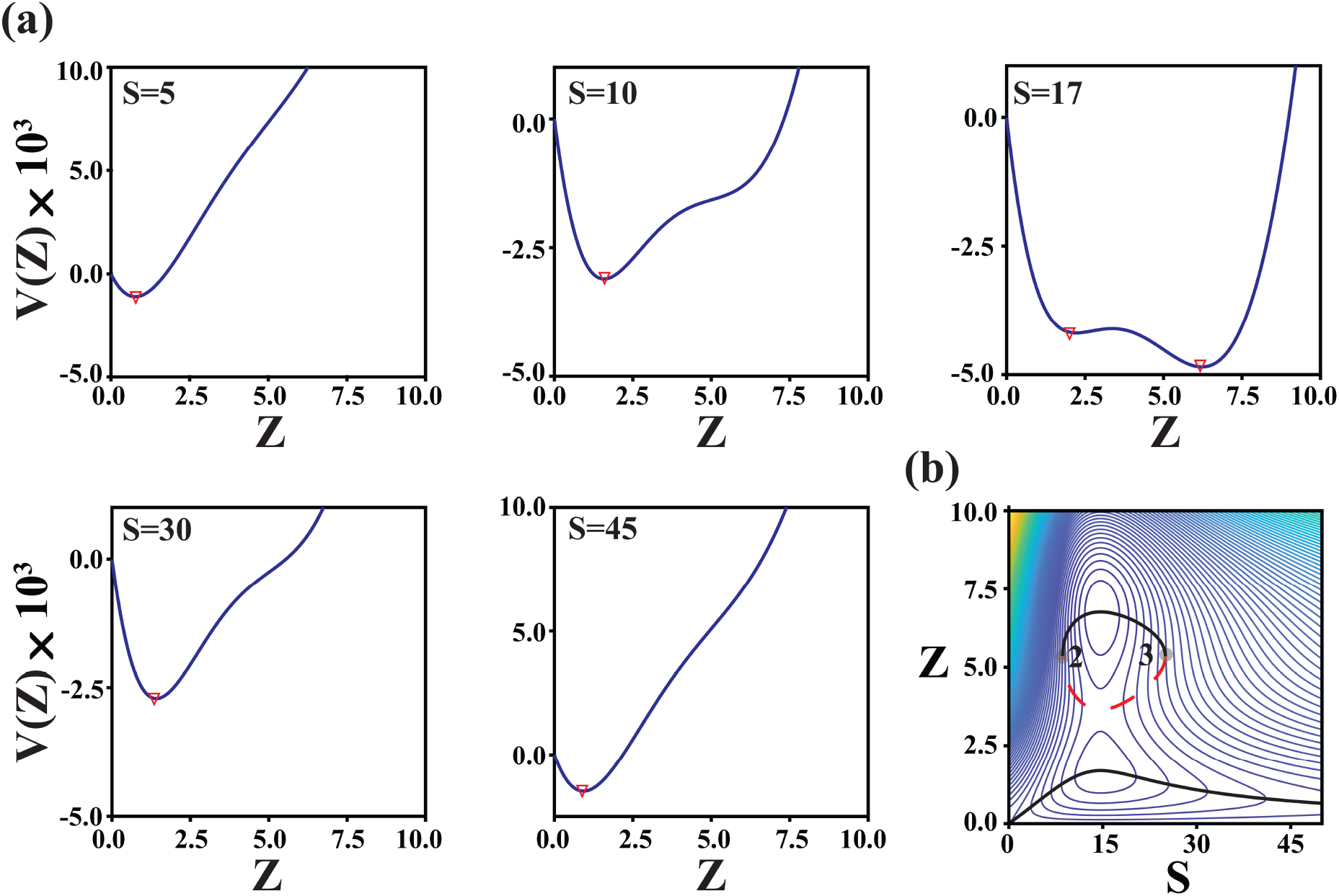
Alteration in the auto-positive regulation leads to Isola bifurcation. **(a)** The plot of potential as a function of *Z* for different values of *S*. **(b)** Energy-based bifurcation diagram showing Isola bifurcation for Network-1. **(c)** Eigenvalue-based bifurcation overlaid over the contour of the potential energy landscape.

We obtain Isola bifurcations for Network-2 (**Fig.S2**) and Network-3 (**Fig.S3**) as well by performing a similar analysis. This indicates that these biological networks have the potential to generate both mushroom and Isola bifurcations under specific conditions.

### Interplay of positive feedback and incoherence decide the nature of bifurcation

However, the features of these networks, which eventually dictate the nature of these bifurcations remain unidentified. Herein, we employed a 2-p bifurcation analysis to explore this issue. The grey line at *k_sZ_* = 0.055 (for which we obtain mushroom bifurcation for network-1 (**Fig.2c**)) in the 2-p bifurcation diagram (**Fig.4a**) of the positive feedback regulatory parameter (*k_sZ_*) and signal strength (*S*) for network-1 reflects that higher degree of positive feedback will lead to greater stabilization of the mushroom bifurcation. However, **Fig.4a** exhibits that any reduced value of *k_sZ_* within the orange-shaded region will lead to an Isola bifurcation, and a further reduction in *k_sZ_* will produce a monostable situation (**Fig.S4b**).

**Fig.4:**
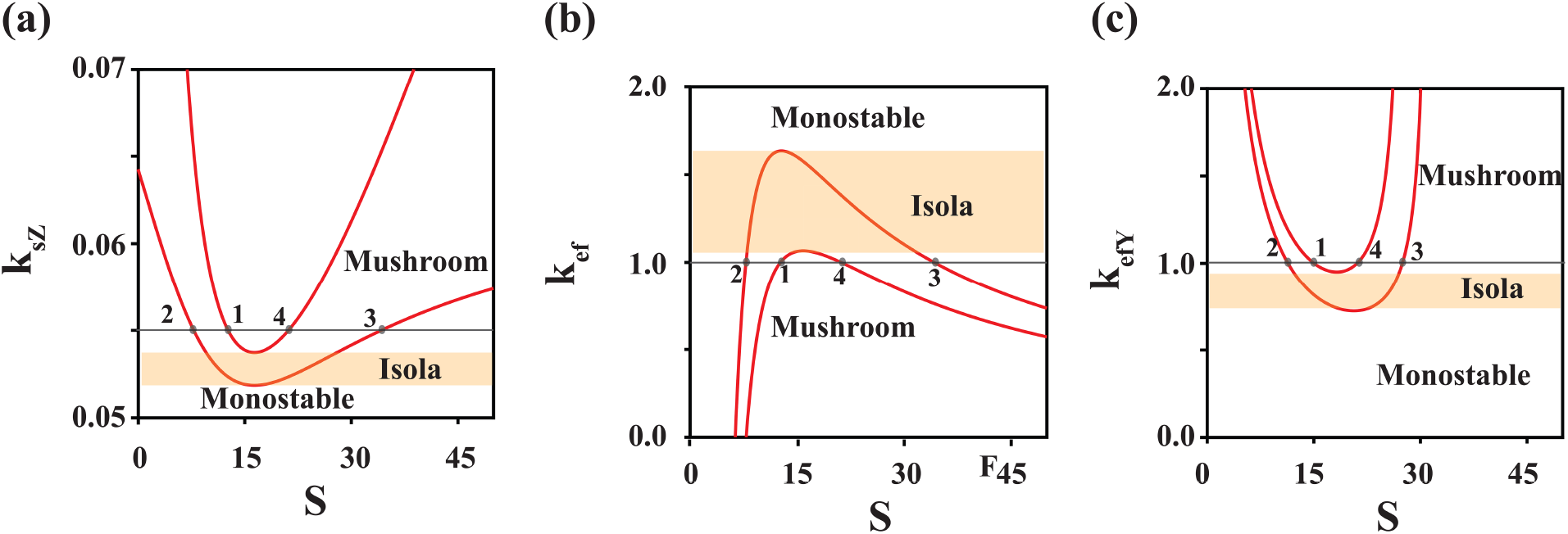
2-parameter bifurcation analysis elucidates the effect of auto positive regulation and incoherence in attaining the Mushroom and Isola bifurcations. Effect of **(a)** auto positive regulation (*k_sZ_ vs S*), and **(b)** incoherence (*k_ef_ vs S*) in obtaining mushroom bifurcation for Network-1. **(c)** Role of incoherence (*k_efY_ vs S*) for Network-3 in achieving mushroom bifurcation.

On the contrary, for network-1, elevating only the level of incoherence (for the node *Y* ⊣ *Z*, (*k_ef_*)) creates an Isola bifurcation scenario (**Fig.4b**) for a range of values for *k_ef_* depicted by the orange-shaded region. While increasing the extent of this negative regulation produces a monostable system (**Fig.S4c**), lowering the same will first maintain the system in the mushroom region for a specific range of *k_ef_* and further Decrease in *k_ef_* will generate a simple bi-stable system (**Fig.4b**). Importantly, the nature of incoherence in the network can be alternatively tuned by altering the parameters associated with the other two nodes (from *X* → *Z*, (*k*_*sZ*1_) and *X* → *Y*, (*k_sY_*)) as well. The respective 2-p bifurcation diagrams reveal that *k_sY_* influences (**Fig.S5a**) the dynamics in a similar manner as that of *k_ef_*, but raising the *k*_*sZ*1_ (**Fig.S5b**) will stabilize the mushroom bifurcation as observed in the case of the positive feedback-related parameter (**Fig.4a**).

We have performed similar 2-p bifurcation analyses for network-2 and 3 (**Fig.1**), which are structurally quite similar to each other. For both these networks, the 2-p bifurcations of *k_sZ_* (extent of positive feedback) as a function of *S* (**Fig.S6a** and **Fig.S7a**) remain qualitatively identical with network-1, as we maintain the same parameters related to the positive feedback interactions in the Hill term representing the auto-positive regulation of *Z*. Fascinatingly, for these three networks, the saddle nodes (SN_1_, SN_2_ SN_3_, and SN_4_) appeared at the same *Z* values for different signal strength *S* (**Table-S3**). This speculates that the auto-positive regulation over *Z* determines the positions of the saddle nodes over the potential energy surfaces, but how and when to reach and exit those saddle points are dictated by the incoherence of the respective networks.

For network-2 and 3, the 2-p bifurcation depicting the variation of extent of incoherence *k_ef_* (the *X* ⊣ *Z* node) as a function of *S* showed similarity with **Fig.4b**, where an increase in *k_ef_* produced isola bifurcation (**Fig.S6b** and **Fig.S7b**). The incoherence from the other two nodes (*X* → *Y* → *Z* (network-2), or *X* ⊣ *Y* ⊣ *Z*, (network-3)) is eventually activating the positive feedback regulation of *Z*. Thus, for network-2, the 2-p bifurcations of *k_sY_* (*Z* → *Y* node) (**Fig.S6c**) and *k*_*sZ*1_ (*Y* → *Z* node) (**Fig.S6d**) with *S* indicates that increasing either one of these two parameters will strengthen the mushroom bifurcation. However, for network-3, the 2-p bifurcation diagram (**Fig.4c**) between the incoherence parameter *k_efY_* (*X* ⊣ *Y* node) with *S* reveals that one can further stabilize the mushroom bifurcation by increasing *k_efY_*. Our investigation thus demonstrates that these three networks are capable of producing Mushroom or Isola bifurcations due to a delicate balance between incoherence of the networks and the extent of positive feedback regulation.

### Network-4 has the capability of producing inverted Mushroom and Isola bifurcations

Till now, we have not discussed network-4, which has a striking similarity with network-1. Interestingly, the potential landscape analysis (**Fig.S8a**) for the model associated with network-4 brings out that at a lower value of *S* (*S* = 5), the system remains in a stable local minimum corresponding to a high value of *Z*. This was quite in contrast with the result observed for network-1 at the same *S* value (**Fig.2a**). Systematically increasing the level of *S* further indicated the existence of an inverted mushroom type bifurcation for network-4 (**Fig.S8a-b**). This was validated by the eigenvalue-based 1-p bifurcation (**Fig.5a**) under the same parametric condition. To investigate what caused this inverted mushroom bifurcation, we resort to a 2-p bifurcation analysis. For network-4, we found that moderately increasing the extent of positive feedback (*k_sZ_*) (orange shaded region, **Fig.5b**) will lead to an inverted Isola bifurcation (**Fig.5c**). Increasing *k_sZ_* too much (**Fig.S8c**), or decreasing it below a definite range (**Fig.S8d**) will create a monostable system. For a moderate range of *k_sZ_*, the system will exhibit mushroom bifurcation. This kind of dynamical feature has been caused by the inverted nature of 2-p bifurcation of the *k_sZ_* as a function of *S* (**Fig.5b**) for network-4 in comparison to the other networks (1-3) (**Fig.4a**).

**Fig.5:**
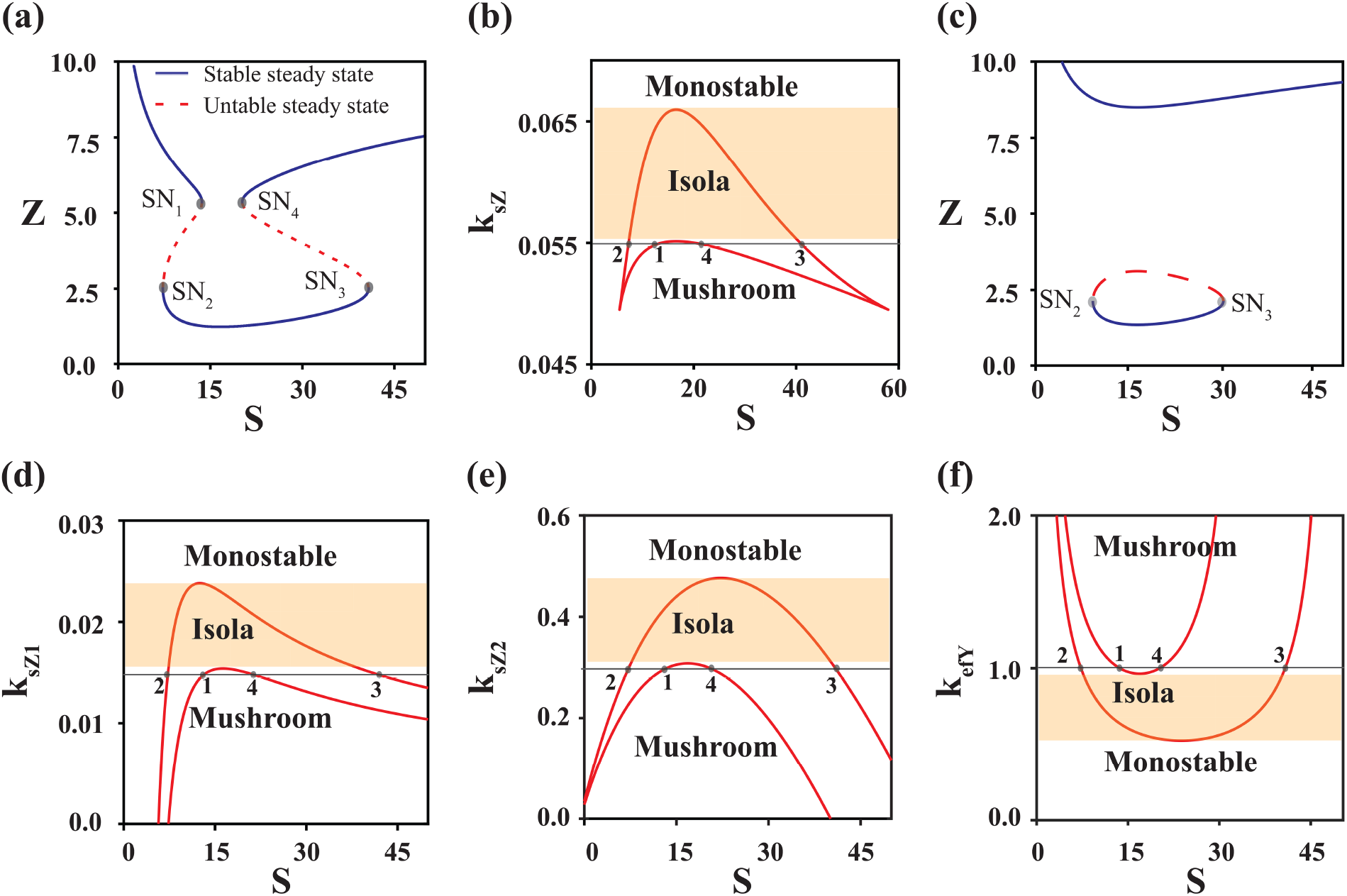
Network-4 is capable of showing inverted Mushroom and Isola bifurcations. **(a)** Eigenvalue-based bifurcation of *Z* vs *S* demonstrates inverted mushroom bifurcation. **(b)** Significance of auto-activation in altering the reverse mushroom bifurcation. **(c)** Inverted Isola bifurcation obtained for *k_sZ_* = 0.06. Influence of positive regulations **(d)** *k*_*sZ*1_ vs *S* and **(e)** *k*_*sZ*2_ vs *S*, and **(f)** negative regulation (*k_efY_* vs *S*) in achieving inverted Mushroom and Isola bifurcations.

Intriguingly, we have kept the same values for the positive feedback related parameters (*k_sZ_* and *k_mZ_*) for network-4 (like network-1), but the order in which the saddle nodes appeared in the 1-p bifurcation (**Fig.5a**) has changed keeping the positions of the saddle nodes intact (**Table S3**). At this point, a careful inspection of the 2-p bifurcations (**Fig.5d-f**) of the incoherence related parameters (*k*_*sZ*1_ (*Z* → *Z* node), *k*_*sZ*2_ (*Y* → *Z* node) and *k_efY_* (*X* ⊣ *Y* node)) with *S* disclose how the inverted Mushroom or Isola bifurcations appear dynamically for network-4. **Fig.4d** and **Fig.4e** ascertain that the positive regulation of *Z* either via *Y* or *X* must remain below a certain threshold level to produce a mushroom bifurcation, and choice of the *k*_*sZ*2_ and *k*_*sZ*1_ (grey lines, **Fig.4e** and **Fig.4d**) insist that *Y* must activate *Z* at a greater extent than *X*. Additionally, **Fig.4f** elucidates that *X* must inhibit *Y* quite strongly to keep the system in the mushroom bifurcation domain. This reiterates the fact that a critical balance of incoherence and the extent of positive feedback regulation determines the ultimate dynamical nature of these networks.

## Conclusion

In this article, we showed that incoherent feedforward loops in combination with an auto-positive feedback motif are sufficient to generate Mushroom and Isola bifurcations. We proposed a set of simple mathematical models for these minimal networks (**Table-S1**), and using the Waddington potential landscape theory and bifurcation analysis, we demonstrated the dynamical diversities of these networks (**Fig.2-5**). Furthermore, we decoded the decisive features of these network motifs that allowed them to generate these complex bifurcations. By employing a 2-p bifurcation study (**Fig.4-5** and **Fig.S5-7**), we investigated the network features that control the existence of a Mushroom (Isola) or a reverse Mushroom (reverse Isola) kind of dynamics. Our analysis revealed that for all the networks (**Fig.1**), the extent of incoherence can be altered to modify the appearance of the bi-stable dynamics due to the auto-positive feedback motif, which eventually diversifies the steady-state dynamics of *Z*.

For network-1, the initial bi-stable activation of (*Z*) was caused by the quick activation of the *X* → *Z* node in comparison to *X* → *Y* node (**Fig.S5**). However, for higher values of *S,* the *X* → *Y* ⊣ *Z* node governed the dynamics (**Fig.3b**) to generate the reverse bi-stable switch in *Z*. In contrast, for network-2, the *X* → *Y* → *Z* node initiated the auto-positive activation of *Z*, and the *X* ⊣ *Z* node influenced the dynamics for higher values of *S* to trigger the mushroom bifurcation (**Fig.S6**). For network-3, it is even more intriguing with all the negative regulatory nodes. The 2-p bifurcations suggest that the first bi-stable activation of *Z* has happened due to the *X* ⊣ *Y* ⊣ *Z* node, where a higher extent of *X* ⊣ *Y* maintains *Y* at a lower level, which allows the initial auto-activation of *Z* (**Fig.S7**). For higher values of *S*, finally, the *X* ⊣ *Z* node dominates the dynamics and brings down *Z* to produce the mushroom bifurcation. Thus, our results indicate that delaying the effect of the direct inhibitory node to *Z* (for network-1-3) is the main reason to produce the Mushroom and Isola bifurcations. However, for network-4, both the direct nodes to *Z* (*X* → *Z* and *Y* → *Z*) are activating it. Our study indicates that if the activation of *Z* due to *Y* → *Z* node is greater (**Fig.5d**) than the activation caused by the *X* → *Z* (**Fig.5e**), and the extent of incoherence due to *X* ⊣ *Y* is higher for the initial level of *S* (**Fig.5f**), then we can have a reverse mushroom bifurcation (**Fig.5a**). This further reasserts that by adjusting the incoherence of these networks, it is possible to observe complex bifurcations in the dynamical systems.

Can we obtain a normal mushroom bifurcation for network-4? In principle, it is possible, as this network is structurally close to network-1. However, the closed cusp-like feature in the 2-p bifurcation of *k_sZ_* vs. *S* (**Fig.5b**) implies that for this simple model (**Table-S1**), under the chosen parametric condition, it is not possible to have a normal mushroom bifurcation. Similarly, reverse Mushroom or reverse Isola bifurcations cannot be observed for the other 3 network models. Even modifying the non-linearities associated with the model equations (**Fig.S9**) does not change this conclusion. This does not mean that these networks are incapable of producing such bifurcations, but with our simple models, it is difficult to generate such scenarios. In summary, our analysis uncovers a set of four minimal network structures that are inherently capable of producing Mushroom and Isola bifurcations. In the future, our findings will allow quick identifications of network motifs that govern the underlying dynamics of complex biological networks by creating these complex bifurcations.

## Data Availability Statement

The Matlab and XPPAUT ODE files related to this work are available at the following link: https://github.com/amitava-giri/Mushroom_bifurcation.git

## Acknowledgments

Thanks are due to UGC for providing the UGC-CSIR-JRF fellowship (Ref. No: 19-06/2016(i)EU-V) to (AG). This work is supported by the funding agency **SERB, India** (Grant no. **CRG/2019/002640** and Grant no. **MTR/2020/000261**).

