## Supplementary material for "Incoherent modulation of bi-stable dynamics orchestrates the Mushroom and Isola bifurcations": Supplementary_materials_bioxriv.pdf

**Table-S1:** Four candidate composite network motifs and the corresponding governing dynamical equations for the networks.

| Network motif | Mathematical model | |
|---|---|---|
| 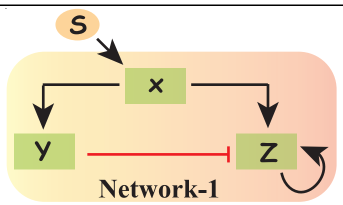  | $\frac{dZ}{dt} = J_{0Z} + \frac{k_{sZ} \times Z^2}{k_{mZ}^2 + Z^2} + \frac{k_{sZ1} \times X}{k_{mZ1} + k_{ef} \times Y^n} - d_Z \times Z$ | 1 |
| | $\frac{dY}{dt} = J_{0Y} + k_{sY} \times X - d_Y \times Y$ | 2 |
| | $\frac{dX}{dt} = J_{0X} + k_{sX} \times S - d_X \times X$ | 3 |
| 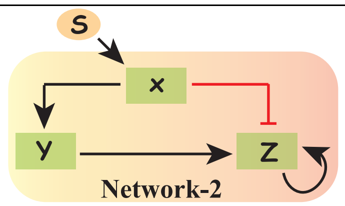  | $\frac{dZ}{dt} = J_{0Z} + \frac{k_{sZ} \times Z^2}{k_{mZ}^2 + Z^2} + \frac{k_{sZ1} \times Y}{k_{mZ1} + k_{ef} \times X^n} - d_Z \times Z$ | 4 |
| | $\frac{dY}{dt} = J_{0Y} + k_{sY} \times X - d_Y \times Y$ | 5 |
| | $\frac{dX}{dt} = J_{0X} + k_{sX} \times S - d_X \times X$ | 6 |
| 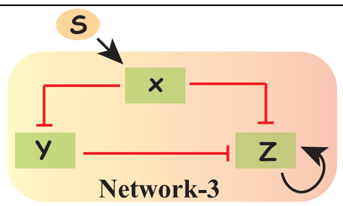  | $\frac{dZ}{dt} = J_{0Z} + \frac{k_{sZ} \times Z^2}{k_{mZ}^2 + Z^2} + \frac{k_{sZ1}}{k_{mZ1} + k_{ef1} \times X^n + k_{ef2} \times Y^n} - d_Z \times Z$ | 7 |
| | $\frac{dY}{dt} = J_{0Y} + \frac{k_{sY}}{k_{mY} + k_{efY} \times X^{n1}} - d_Y \times Y$ | 8 |
| | $\frac{dX}{dt} = J_{0X} + k_{sX} \times S - d_X \times X$ | 9 |
| | $\frac{dZ}{dt} = J_{0Z} + \frac{k_{sZ} \times Z^2}{k_{mZ}^2 + Z^2} + \frac{k_{sZ1} \times X + k_{sZ2} \times Y}{k_{mZ1} + X + Y} - d_Z \times Z$ | 10 |

| | | |
|---|---|---|
| 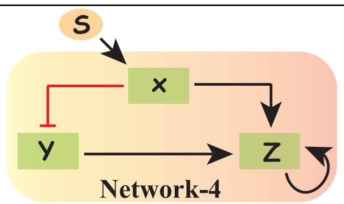 | $\frac{dY}{dt} = J_{0Y} + \frac{k_{sY}}{k_{mY} + k_{efY} \times X^{n1}} - d_Y \times Y$ | 11 |
| | $\frac{dX}{dt} = J_{0X} + k_{sX} \times S - d_X \times X$ | 12 |

**Table S2**: Description and values of each parameter used in the mathematical model.

| Symbol | Description | Value | Unit |
|---|---|---|---|
| **Network 1** | | | $c = conc.$ $t = time$ |
| $J_{0X}$ | Basal synthesis of X | $10^{-6}$ | $c\,t^{-1}$ |
| $k_{sX}$ | Activation rate of X by the external signal | 0.001 | $t^{-1}$ |
| $d_X$ | Degradation of X | 0.003 | $t^{-1}$ |
| $J_{0Y}$ | Basal synthesis of Y | $5 \times 10^{-5}$ | $c\,t^{-1}$ |
| $k_{sY}$ | Activation rate of Y by X | 0.0011 | $t^{-1}$ |
| $d_Y$ | Degradation rate of Y | 0.003 | $t^{-1}$ |
| $J_{0Z}$ | Basal synthesis of Z | $5 \times 10^{-5}$ | $c\,t^{-1}$ |
| $k_{sZ}$ | Auto activation rate of Z | 0.055 | $c\,t^{-1}$ |
| $k_{mZ}$ | Michaelis constant of auto activation of Z | 6.5 | $c$ |
| $k_{sZ1}$ | Activation rate of Z by X | 0.008 | $c^2\,t^{-1}$ |
| $k_{mZ1}$ | Michaelis type constant | 4.0 | $c^2$ |
| $k_{ef}$ | Repression rate of Z by Y | 1.0 | - |
| $n$ | Hill coefficients corresponds to the repression by Y to Z | 2.0 | - |
| $d_Z$ | Degradation constant of Z | 0.00495 | $t^{-1}$ |
| **Network 2** | | | |
| $J_{0X}$ | Basal synthesis of X | $10^{-6}$ | $c\,t^{-1}$ |
| $k_{sX}$ | Activation rate of X by the external signal | 0.0002 | $t^{-1}$ |
| $d_X$ | Degradation of X | 0.003 | $t^{-1}$ |

| | | | |
|---|---|---|---|
| $\boldsymbol{J_{0Y}}$ | Basal synthesis of Y | $5 \times 10^{-5}$ | $c\, t^{-1}$ |
| $\boldsymbol{k_{sY}}$ | Activation rate of Y by X | 0.024 | $t^{-1}$ |
| $\boldsymbol{d_Y}$ | Degradation rate of Y | 0.003 | $t^{-1}$ |
| $\boldsymbol{J_{0Z}}$ | Basal synthesis of Z | $5 \times 10^{-5}$ | $c\, t^{-1}$ |
| $\boldsymbol{k_{sZ}}$ | Auto activation rate of Z | 0.055 | $c\, t^{-1}$ |
| $\boldsymbol{k_{mZ}}$ | Michaelis constant of auto activation of Z | 6.5 | $c$ |
| $\boldsymbol{k_{sZ1}}$ | Activation rate of Z by Y | 0.002 | $c^2\, t^{-1}$ |
| $\boldsymbol{k_{mZ1}}$ | Michaelis type constant | 2.25 | $c^2$ |
| $\boldsymbol{k_{ef}}$ | Repression rate of Z by X | 1.0 | - |
| $\boldsymbol{n}$ | Hill coefficients corresponds to the repression by X to Z | 2.0 | - |
| $\boldsymbol{d_Z}$ | Degradation constant of Z | 0.00495 | $t^{-1}$ |
| **Network 3** | | | |
| $\boldsymbol{J_{0X}}$ | Basal synthesis of X | $10^{-6}$ | $c\, t^{-1}$ |
| $\boldsymbol{k_{sX}}$ | Activation rate of X by the external signal | 0.0001 | $t^{-1}$ |
| $\boldsymbol{d_X}$ | Degradation of X | 0.003 | $t^{-1}$ |
| $\boldsymbol{J_{0Y}}$ | Basal synthesis of Y | 0.00025 | $c\, t^{-1}$ |
| $\boldsymbol{k_{sY}}$ | Synthesis rate of Y | 0.00027 | $c^2\, t^{-1}$ |
| $\boldsymbol{k_{efY}}$ | Repression strength by X on Y | 1.0 | - |
| $\boldsymbol{k_{mY}}$ | Michaelis type constant related to inhibition of Y by X | 0.001 | $c$ |
| $\boldsymbol{d_Y}$ | Degradation rate of Y | 0.003 | $t^{-1}$ |
| $\boldsymbol{J_{0Z}}$ | Basal synthesis of Z | $2.5 \times 10^{-5}$ | $c\, t^{-1}$ |
| $\boldsymbol{k_{sZ}}$ | Auto activation rate of Z | 0.055 | $c\, t^{-1}$ |
| $\boldsymbol{k_{mZ}}$ | Michaelis constant of auto activation of Z | 6.5 | $c$ |
| $\boldsymbol{k_{sZ1}}$ | Synthesis rate of Z | 0.0005 | $c^3\, t^{-1}$ |
| $\boldsymbol{k_{ef1}}$ | Repression strength by X | 0.1 | - |
| $\boldsymbol{k_{ef2}}$ | Repression strength by X | 1.0 | - |

| | | | |
|---|---|---|---|
| $n$ | Hill coefficient corresponding to the repression by X and Y | 2.0 | - |
| $n1$ | Hill coefficients corresponding to the repression by X on Y | 1.0 | - |
| $k_{mZ1}$ | Michaelis type constant related to repression of Z by X and Y | 0.001 | $c^2$ |
| $d_Z$ | Degradation constant of Z | 0.00495 | $t^{-1}$ |
| **Network 4** | | | |
| $J_{0X}$ | Basal synthesis of X | $10^{-6}$ | $c\,t^{-1}$ |
| $k_{sX}$ | Activation rate of X by the external signal | 0.00025 | $t^{-1}$ |
| $d_X$ | Degradation of X | 0.003 | $t^{-1}$ |
| $J_{0Y}$ | Basal synthesis of Y | $5 \times 10^{-5}$ | $c\,t^{-1}$ |
| $k_{sY}$ | Synthesis rate of Y | 0.000225 | $c^2\,t^{-1}$ |
| $k_{efY}$ | Repression strength by X on Y | 1.0 | - |
| $n1$ | Hill coefficients corresponding to the repression by X on Y | 1.0 | - |
| $k_{mY}$ | Michaelis type constant related to inhibition of Y by X | 0.05 | $c$ |
| $d_Y$ | Degradation rate of Y | 0.003 | $t^{-1}$ |
| $J_{0Z}$ | Basal synthesis of Z | $5 \times 10^{-5}$ | $c\,t^{-1}$ |
| $k_{sZ}$ | Auto activation rate of Z | 0.055 | $c\,t^{-1}$ |
| $k_{mZ}$ | Michaelis constant of auto activation of Z | 6.5 | $c$ |
| $k_{sZ1}$ | Activation rate of Z by X | 0.015 | $c\,t^{-1}$ |
| $k_{sZ2}$ | Activation rate of Z by Y | 0.3 | $c\,t^{-1}$ |
| $k_{mZ1}$ | Michaelis type constant related to activation of Z by X and Y | 8.5 | $c$ |
| $d_Z$ | Degradation constant of Z | 0.00495 | $t^{-1}$ |

**Table S3**: Positions of bifurcation points in all the network structures.

| Network | $S$ | $Z$ | Saddle node bifurcation point |
|---|---|---|---|
| Network 1 | 12.66 | 2.5111 | $SN_1$ |
| | 7.830 | 5.3502 | $SN_2$ |
| | 34.194 | 5.3502 | $SN_3$ |
| | 21.148 | 2.5111 | $SN_4$ |
| Network 2 | 18.419 | 2.5111 | $SN_1$ |
| | 10.992 | 5.3502 | $SN_2$ |
| | 45.865 | 5.3502 | $SN_3$ |
| | 27.395 | 2.5111 | $SN_4$ |
| Network 3 | 15.038 | 2.5111 | $SN_1$ |
| | 11.369 | 5.3502 | $SN_2$ |
| | 27.457 | 5.3502 | $SN_3$ |
| | 21.078 | 2.5111 | $SN_4$ |
| Network 4 | 13.647 | 5.3502 | $SN_1$ |
| | 7.271 | 2.5111 | $SN_2$ |
| | 40.803 | 2.5111 | $SN_3$ |
| | 20.190 | 5.3502 | $SN_4$ |

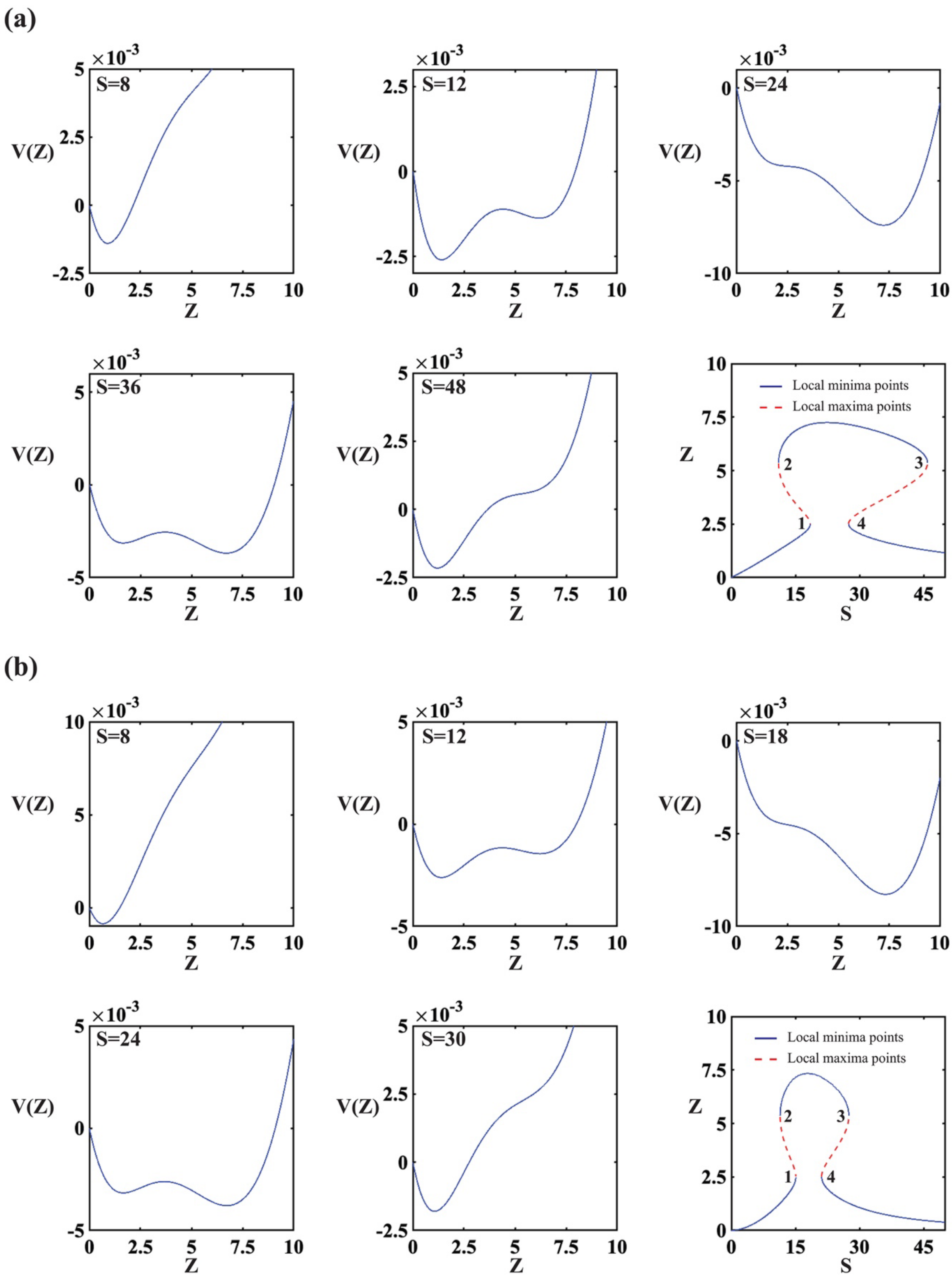

**Fig.S1**: Potential landscape analysis on (**A)** Network 2 and **(B)** Network 3. The last panel of **(A)** and **(B)** is the potential energy based bifurcation diagram of Network 2 and Network 3 respectively. Energy based bifurcation has been constituted by determining the local minima and maxima points. Local minima represents the stable steady state and the local maxima points represent unstable steady state.

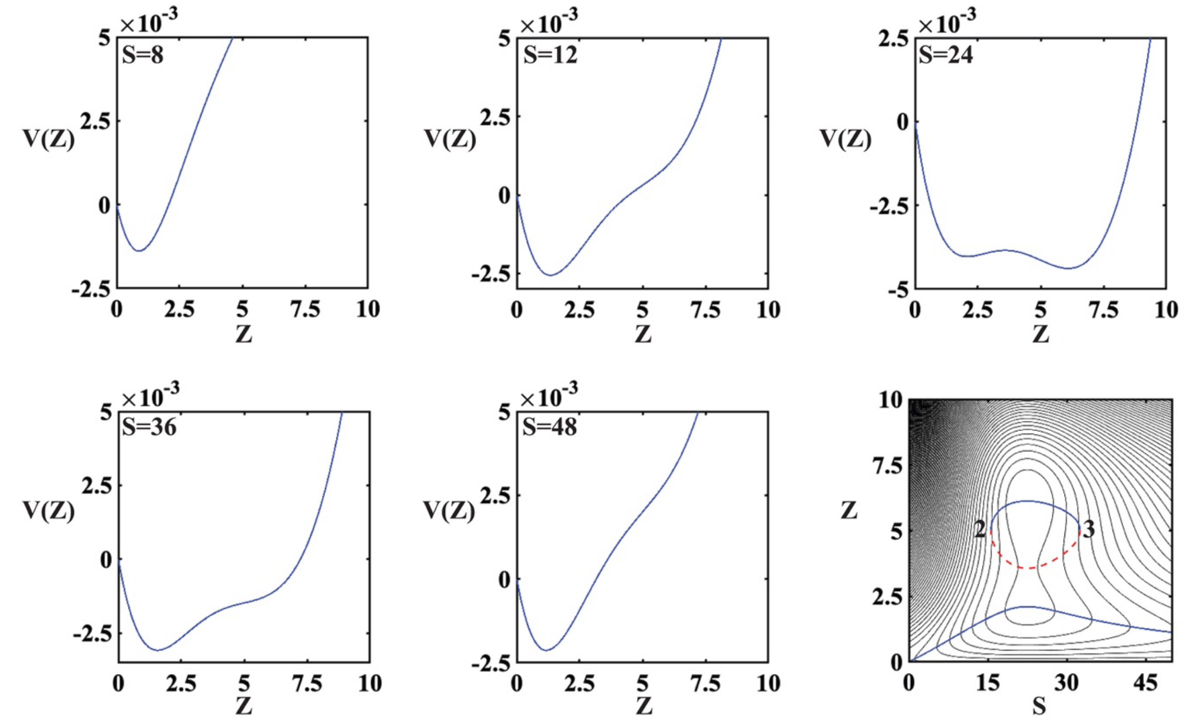

**Fig.S2**: Alteration of auto positive regulation leads to isola bifurcation for Network 2. The plot of potential as a function of Z shows the existence of single local minima for low(S=5,10) and high external signal strength (S=30, 45) but there is two local minima and one local maxima at moderate external signal strength (S=17). In the last panel eigen value based bifurcation is overlaid on the potential energy contour plot.

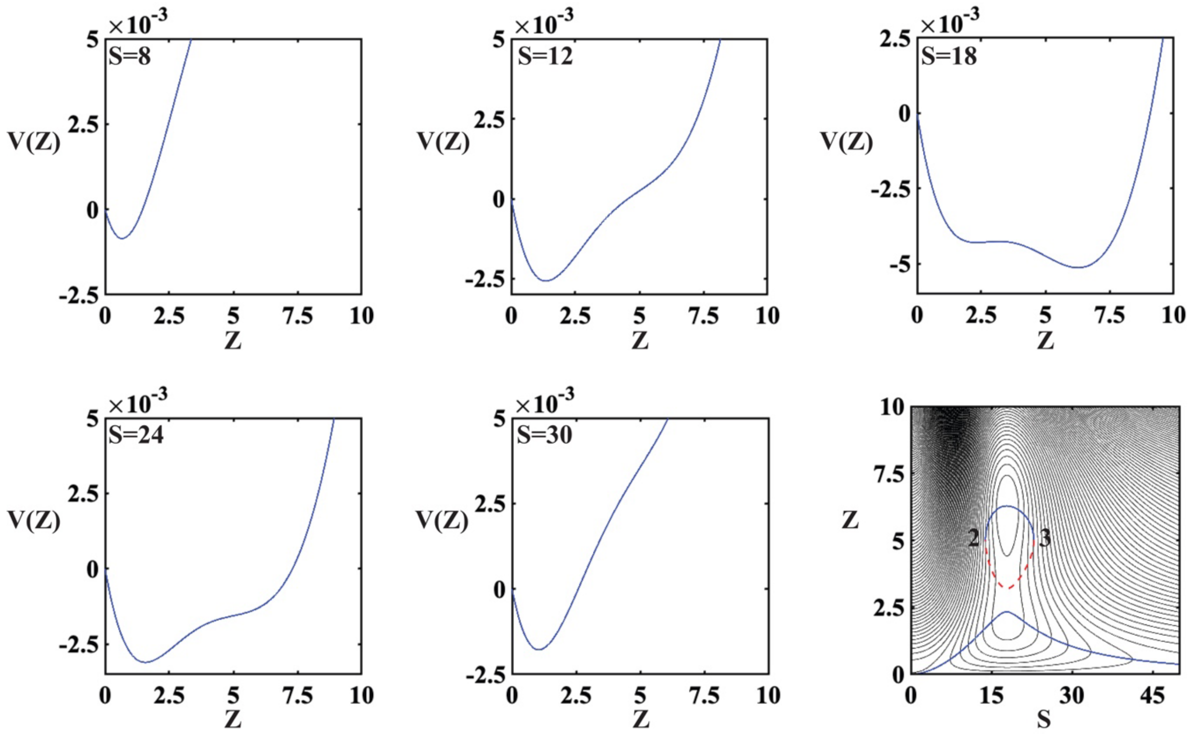

**Fig.S3**: Alteration of auto positive regulation leads to isola bifurcation for Network 3.

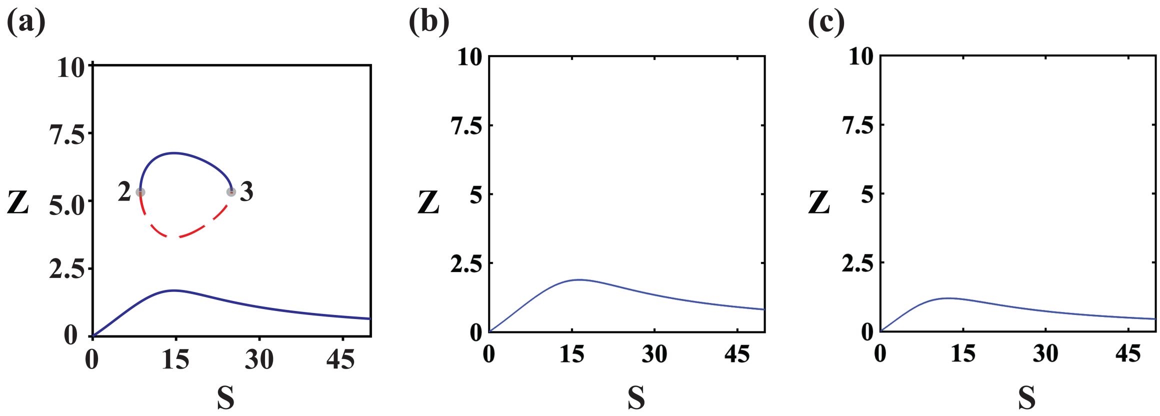

**Fig.S4**: Either **(A)** decrease of extent of auto positive regulation or **(B)** increase the extent of incoherence leads to monostable situation.

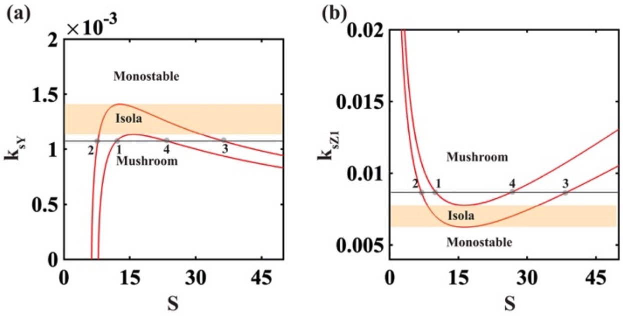

**Fig.S5**: Two parameter bifurcations analysis reveals the role of **(A)** X activating Y node ($k_{sY}$ $vs$ $S$) and **(B)** X activating Z node ($k_{sZ1}$ $vs$ $S$) in achieving mushroom and isola bifurcation features for Network 1.

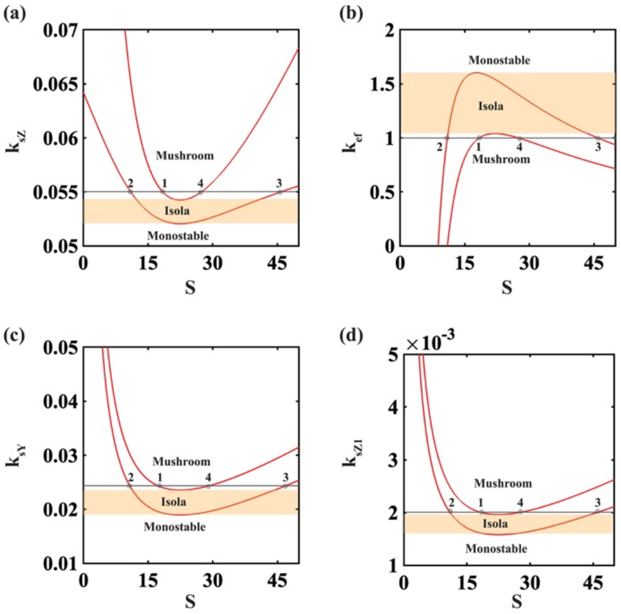

**Fig.S6**: Role of different nodes in achieving mushroom and isola bifurcation for Network 2**. (A)** Auto activation node ($k_{sZ}$ $vs$ $S$) **(B)** X inhibiting Z node ($k_{ef}$ $vs$ $S$) **(C)** X activating Y node ($k_{sY}$ $vs$ $S$) and **(D)** Y activating Z node ($k_{sZ1}$ $vs$ $S$).

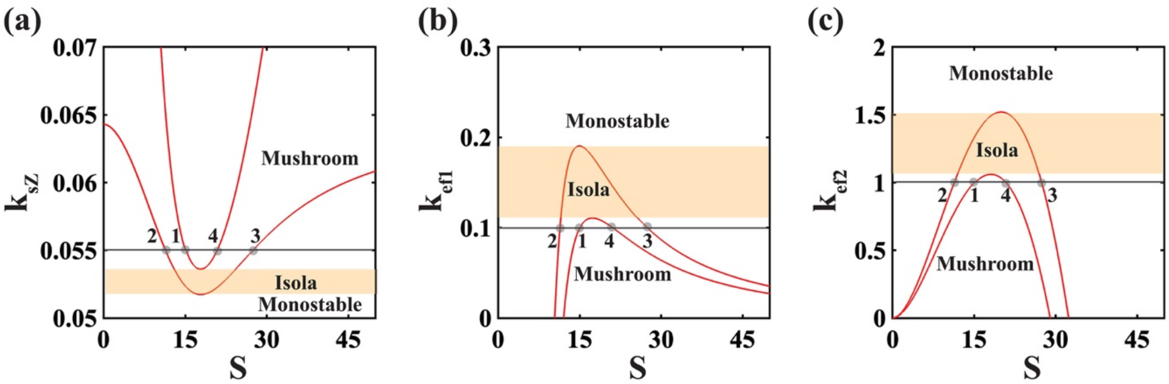

**Fig.S7**: Role of different nodes in achieving mushroom and isola bifurcation for Network 3. **(A)** auto activation node ($k_{sZ}$ $vs$ $S$) **(B)** X inhibiting Z node ($k_{ef1}$ $vs$ $S$) **(C)** Y inhibiting Z node ($k_{ef2}$ $vs$ $S$).

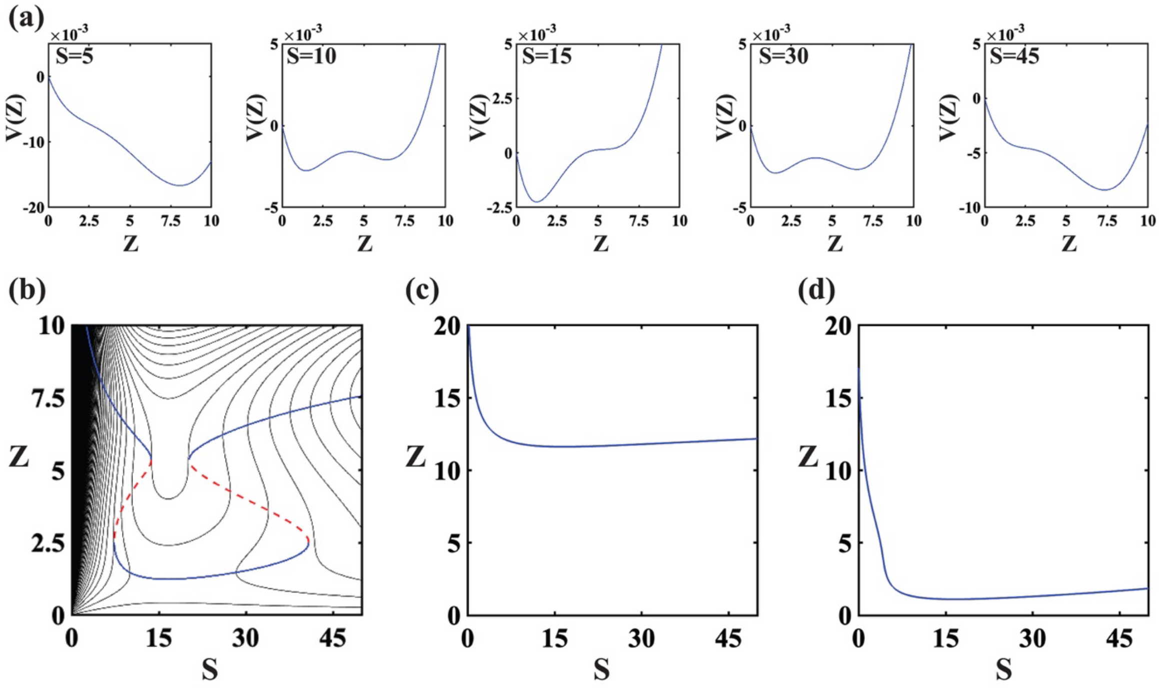

**Fig.S8:** Study of effective potential as a function of Z for different external signal, shows the existence of reverse mushroom type of bifurcation. **(A)** plot of potential as a function of Z. **(B)** Eigen value based bifurcation diagram is overlayed on top of the potential energy contour for Network 4. **(C)** High $k_{sZ}$ value gives monostable scenario. **(d)** Low $k_{sZ}$ value also leads to monostable situation.

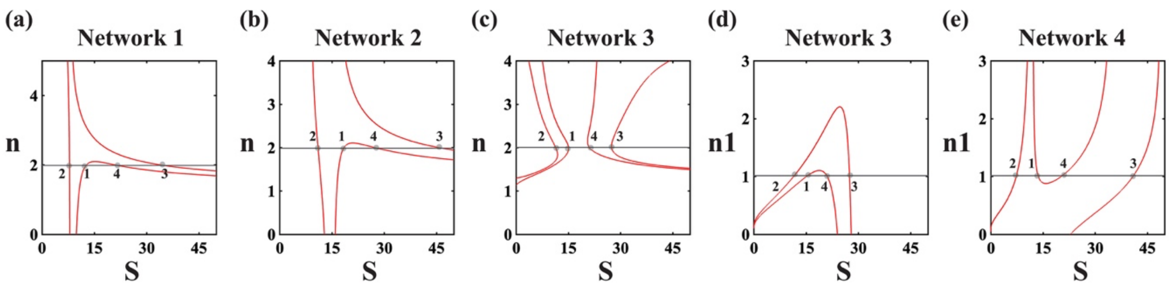

**Fig.S9:** Effect of nonlinearity in the mathematical model. Two parameter bifurcations of Hill coefficient vs external signal are drawn for all the networks.
